# A facile green synthesis approach to silver nanoparticles using calyx from *Abelmoschus esculentus* and its anthelmintic activity

**DOI:** 10.1101/2023.03.20.533571

**Authors:** Rima Majumdar, Pradip Kumar Kar

## Abstract

In recent years, technology pertaining to nanobiomaterials has taken rapid strides, with the development of novel materials having unique properties. Silver nanoparticles (AgNPs) have gained attention among these materials due to their high chemical stability, surface-to-volume ratio, and strong antimicrobial activity. The traditional method for synthesizing AgNPs involves toxic chemicals, which can have negative environmental impacts and pose health risks. Hence, there is a growing need for green synthesis methods for AgNPs that are environmentally friendly and safe for animal and human health. In this study, we explore the green synthesis of AgNPs using calyx from *Abelmoschus esculentus*, also known as okra, as an anthelmintic. *Raillietina* spp. is a common poultry parasite causing significant economic losses to the livestock industry. It is a major cause of ailment and mortality in livestock, deterring the host health. While chemical-based anthelmintic drugs are available, the increasing prevalence of drug-resistant parasite strains has made searching for new and effective treatments imperative. Although ethnomedicine has been promising for treating various diseases, including parasitic infections, nanoparticles have emerged as a viable alternative to traditional anthelmintic curative development. Our study aims at investigating the recent advances in nanomedicine, particularly AgNPs, as anthelmintic agents, which has shown remarkable alterations in the levels of tegumental enzymes, eventually leading to their paralysis and death. We discuss the mechanisms of action of AgNPs against *Raillietina* spp. and highlight the potential benefits of using biosynthesized curatives that interfere with the host-parasite interface to treat parasite-related disorders.

## 1. Introduction

Nanomedicine is a promising field that has the potential to transform the way we approach medical treatments. However, the traditional method for synthesizing AgNPs involves toxic chemicals, which can have negative environmental impacts and pose health risks (Mubayi et al., 2012; Rajeshkumar and Bharath, 2017). To address these concerns, several studies have explored new methods for synthesizing AgNPs that are environment-friendly and safe for human health (Mamillapalli, 2016; Rai et al., 2022). Green synthesis of nanoparticles from natural sources has emerged as an eco-friendly and sustainable approach to producing nanoparticles with various applications in diverse fields. Previous studies have reported AgNP synthesis from diverse plants to explore their antimicrobial properties (Juan Carlos et al., 2020; Ogunsona et al., 2020; Spirescu et al., 2021). Additionally, AgNPs have shown potential for broader spectrum activity and are effective against drug-resistant strains of microorganisms (Namivandi-Zangeneh et al., 2021). *Abelmoschus esculentus*, commonly known as okra, has been used in traditional ethnomedicine in many cultures and passed down through generations to treat various health conditions. Okra pod is believed to have respiratory benefits and is used to treat respiratory ailments such as asthma (Romdhane et al., 2020). It has been conventionally used to treat digestive disorders such as diarrhea and gastrointestinal inflammation (Alves et al., 2018). Several reports have shown its significant effect on treating conditions such as arthritis and joint pain, promoting cardiovascular health through cholesterol-lowering effects, and helping manage blood sugar levels (Elkhalifa et al., 2021; Liu et al., 2021). The okra plant contains several bioactive compounds, including flavonoids, polyphenols, and tannins, which have been reported to exhibit significant antimicrobial activity (Yora et al., 2018). Indeed, okra extract-assisted CoFe_2_O_4_ nanoparticles are proven to have significantly high antimicrobial activity against bacteria and fungal strains (Kombaiah et al., 2018).

In developing countries like India, gastrointestinal helminths lead to substantial health problems in poultry, including weight loss, stunted growth, reduced egg production, and even death in severe cases (Sarba et al., 2019). *Raillietina* is a genus of parasites belonging to the class Cestoda, order Cyclophyllidea, and family Davaineidae, that commonly infects the intestines of birds and mammals, including domestic poultry. Several anthelminthic drugs are used to treat parasitic infestation for *Raillietina* spp. include piperazine, fenbendazole, and levamisole. Piperazine paralyzes the tapeworms, causing them to lose their grip on the gut wall and be expelled from the host’s body (Amemor et al., 2021). The tegument of *Raillietina* performs vital functions such as nutrient absorption and secretion by functioning as a physical barrier against the host immune system (Hrckova et al., 2013). Fenbendazole is a broad-spectrum anthelminthic effective against a wide range of parasitic worms, including *Raillietina* and works by inhibiting the tapeworm’s ability to absorb glucose, causing it to starve and eventually die (Saemi Soudkolaei et al., 2021). Levamisole is another anthelminthic drug commonly used to treat *Raillietina* infections by stimulating the host’s immune system to attack and reduce the tapeworm burden (Gao et al., 2021). However, the prolonged unregulated use of such synthetic anthelmintics has been associated with developing resistance in parasites (Zahedi et al., 2022). Previous studies revealed a range of ethnomedicinal anthelmintics used to treat *Raillietina* spp.; these included plant-based remedies, such as Neem (Azadirachta i*ndica*), Wormwood (*Artemisia annua*), and Aloe vera (*Aloe barbadensis miller*), as well as animal-based remedies, such as bee venom (Ash et al., 2017; Shelke et al., 2020; Verbitskaya and Olechnovich, 2007). Several ethnomedicinal studies confirmed these remedies to treat parasitic infections in rural communities, which exhibited potent anthelmintic activity against *Raillietina* spp. (Giri et al., 2021). Kar et al. previously reported the *in vitro* anthelmintic activity of gold nanoparticles from the fungus *Nigrospora oryzae* on cestode parasites. They observed changes in enzyme activity and disruption of the parasite’s tegument following treatment with gold nanoparticles (Kar et al., 2014).

In the current investigation, we focus on the anthelmintic activity of AgNPs synthesized from *A. esculentus* against *Raillietina* spp., a common intestinal parasite of poultry birds (*Gallus gallus domesticus*). We report the anthelmintic activities of the biosynthesized AgNPs against *Raillietina* spp. using several approaches, including histochemical and ultrastructural studies, which present a promising avenue for developing sustainable and effective anticestodal agents.

## 2. Materials and Methods

### 2.1. Collection of plant material

*Abelmoschus esculentus* was collected from the adjacent agricultural land near the Cooch Behar Panchanan Barma University campus (latitude 26.32213 °N, longitude 89.46015 °E). The taxonomical voucher specimen (Ac-97299) is submitted to the herbarium of the Botanical Survey of India, Eastern Regional Centre, Shillong.

### 2.2. Preparation of plant extract using calyx of *A. esculentus*

The okra pods were washed twice with deionized water, air-dried, chopped and separated from the calyces. Chopped calyces (20 g) were placed in a beaker with 100 ml deionized water and heated in a temperature-controlled water bath for 10 minutes at 90°C. Once cooled down, the extract was filtered (Whatman paper No. 1 filter paper) and kept at a temperature of 4°C until it was needed again.

### 2.3 Phytochemical screening

Preliminary phytochemical screening for principal secondary metabolites of the okra calyces was conducted using the standard qualitative methods with the slightest modifications (Nortjie et al., 2022; Shaikh and Patil, 2020). The extract was investigated for the potential biomolecules associated with reducing silver ions to silver atoms. Phytochemical characterization was performed qualitatively for alkaloids, flavonoids, saponins, glycosides, steroids, tannins, terpenoids and proteins.

### 2.4 Biosynthesis of *A. esculentus* silver nanoparticles (AE-AgNP)

To synthesize AE-AgNPs, a solution of 1 mM AgNO3 (Merck Laboratories, India) was prepared by adding 10 mL of the calyx aqueous extract to a 90 mL aqueous solution, and the mixture was left at room temperature. The color of the solution changed from pale yellow to brown, suggesting AgNP formation owing to the reaction between the extract of AE and silver metal ions. A control set was established by preparing a silver nitrate solution without adding calyx extract, which exhibited no color change. To purify the AE-AgNPs, the extract was removed by centrifugation three times at 15,000 rpm for 20 minutes and washed twice with double-sterilized water.

### 2.5 Characterization studies

Characterization of AgNPs from AE involved analyzing their physical and chemical properties. Initially, the conversion of silver ions in solution through bio-reduction was assessed by the visible alterations in color, which were subsequently tracked using UV-visible (UV-Vis) absorption spectroscopy. The functional groups that stabilized the biosynthesized nanoparticles were determined using Fourier transform infrared spectroscopy (FTIR), while energy-dispersive X-ray (EDX) examination revealed the existence of Ag metal in the sample. The synthesized AgNPs were transformed into a powder through freeze-drying, after which they were subjected to X-ray diffraction (XRD) analysis for crystalline phase identification. Finally, transmission electron microscopy (TEM) imaging was used to envisage the size and structure of the nanoparticles directly.

### 2.6 Collection of parasites and *in vitro* treatments

Freshly sacrificed domestic fowl (*Gallus gallus domesticus* L.) were examined for live mature *Raillietina* spp. These parasites were collected from local abattoirs in Cooch Behar, India and preserved in 0.9% phosphate-buffered saline (PBS) at 37 ± 1°C in an incubator. Control parasites were kept in 0.9% PBS without AE-AgNP at 37 ± 1°C while treatment was performed by directly incubating live worms in varied concentrations (25, 50, 75, 100, 125 µg/ml, 0.9% PBS) of AE-AgNP in separate Petri dishes. Genistein (GEN) was used as a broad-spectrum reference drug at 125 µg/ml of PBS. Six replicates were prepared for each pair of incubation conditions, and the times required to reach the paralytic state and death were recorded. By removing the treated parasites from the test medium and immersing them in slightly warm water, parasite mortality was verified. Once all traces of movement had stopped, the times taken for paralysis and death were recorded. The treated and control parasites were primed further for histochemical localization and scanning electron microscopic studies.

### 2.7 Scanning electron microscopic studies

The paralyzed parasites were stored in neutral buffered formalin (10%) for 24 hours at 4 °C for fixation. Those parasites were rinsed with PBS and dehydrated using gradually increasing acetone concentrations until completely dry. The specimens were then subjected to critical-point drying, and the resulting material was coated with platinum in an ion sputter (JFC-1100, JEOL). Finally, the parasite specimens were visualized under a scanning electron microscope (EVO 18, Zeiss) at an accelerating voltage of 10-15 kV.

### 2.8 Histochemical localization of enzymes

The histochemical investigation was conducted on tegumental enzymes using frozen sections prepared and cut to 10-12 µm in a cryostat (CM 3050S, Leica). The modified lead nitrate method detected acid phosphatase (AcPase) activity in specimens fixed with cold formol calcium (Pearse, 1968). The areas of AcPase activity were indicated by a brownish precipitate on the tegumental sections. Alkaline phosphatase (AlkPase) activity was detected at room temperature (17-20 °C) using the modified coupling azo-dye method. The calcium method of Pearse was followed to locate adenosine triphosphatase (ATPase) activity, using adenosine triphosphate (ATP) as the substrate, and enzyme activity was identified by observing blackish-brown deposits (Pearse, 1968). The Wachstein and Meisel lead method detected 5’-Nucleotidase (5’-Nu) activity, with adenosine monophosphate as the substrate (Wachstein and Meisel, 1957).

## 3. Results and Discussion

### 3.1 Phytochemical screening of *A. esculentus*

The data from the preliminary screening of *A. esculentus* calyx extracts indicated the presence of tannins, alkaloids, saponins, terpenoids, flavonoids, glycosides and proteins. These compounds may actively reduce silver ions to nanoparticles. In Table 1, the presence of phytomolecules is represented by a positive sign (+), their abundance by a double positive sign (++) and their absence by a negative sign (−). The phytochemical results showing the abundance of secondary metabolites are consistent with the findings of Refs. (Nortjie et al., 2022; Shaikh and Patil, 2020).

**Table 1.** Preliminary qualitative screening of the phytochemicals in the aqueous extracts of *A. esculentus* calyces

| Phytochemical constituents | Tests | Aqueous extract |
| --- | --- | --- |
| Alkaloids | Mayer's test | + |
| Saponins | Frothing test | + |
| Glycosides | Keller-Killiani test, Legal's test | — |
| Tannins | Ferric chloride test | ++ |
| Flavonoids | Alkaline reagent test | ++ |
| Terpenoids | Concentrated H<sub>2</sub>SO<sub>4</sub> test | ++ |
| Steroids | Liebermann-Burchard's test | — |
| Proteins | Millon's reagent, Burette's test | ++ |

**Table 2.** *In vitro* efficacy of AE-AgNPs and reference drug Genistein on *Raillietina spp*.

| Test material | Concentration ($\mu\text{g/ml}$ PBS) | Paralysis (h) | Death (h) |
| --- | --- | --- | --- |
| Control | – | $72 \pm 0.06$ | |
| AE-AgNP | 25 | $1.57 \pm 0.16$ | $2.54 \pm 0.14$ |
| | 50 | $1.35 \pm 0.08$ | $2.32 \pm 0.09$ |
| | 75 | $1.06 \pm 0.04$ | $1.54 \pm 0.08$ |
| | 100 | $0.58 \pm 0.08$ | $1.46 \pm 0.04$ |
| | 125 | $0.54 \pm 0.02$ | $1.29 \pm 0.02$ |
| Genistein | 125 | $0.53 \pm 0.02$ | $1.26 \pm 0.03$ |

### 3.2 Characterization of biosynthesized AgNPs

#### 3.2.1 UV-Visible Spectrophotometric Analysis

The formation of AgNPs was assessed by measuring the UV-Vis spectrum of the medium in the range of wavelength from 200 to 700 nm (Fig. 1**A**). AgNPs exhibit a characteristic absorption peak around 425 nm for AgNPs in the UV-Vis spectrum, similar to previously reported results (Awwad and Salem, 2012) and owes to the plasmon oscillation of silver to generate an electric field, known as surface plasmon resonance (SPR).

**Figure 1.**
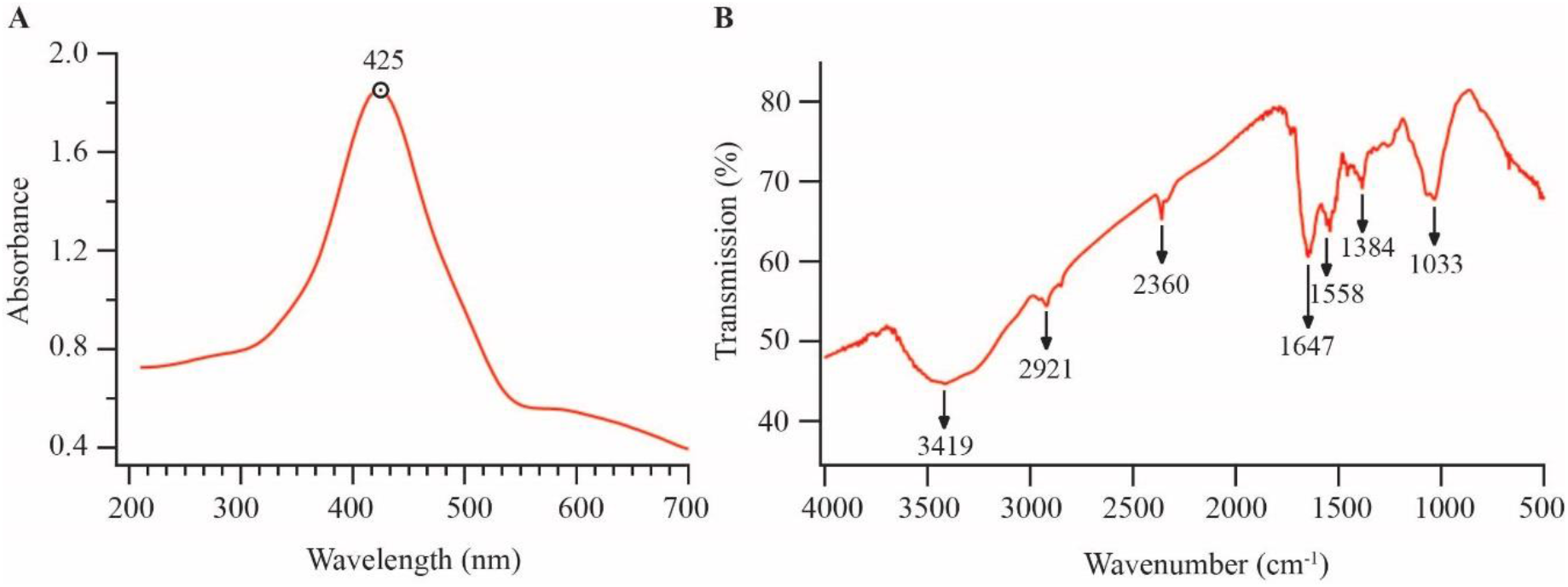
**A**. UV-Vis spectra of AgNPs synthesized from *A. esculentus* extract; **B**. FTIR spectrum of synthesized AgNPs from aqueous extract of *A. esculentus*.

**Figure 2.**
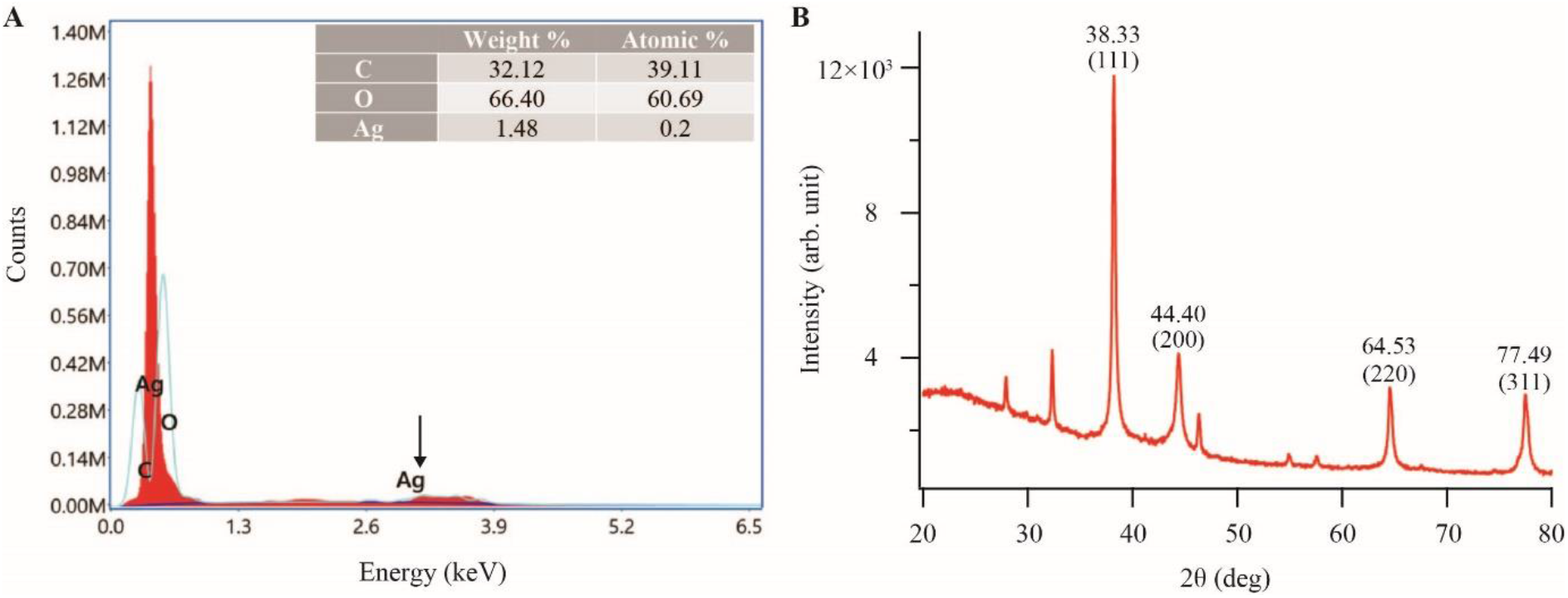
Energy dispersive X-ray spectroscopic spectrum **(A)** and X-ray diffraction pattern **(B)** of the biosynthesized AE-AgNPs.

#### 3.2.2 FTIR Analysis

The FTIR spectrum of dried AgNPs was analyzed to determine the phytochemical constituents responsible for the capping of AgNPs by attributing the absorption bands with their corresponding compounds. The distinct peaks observed from the calyx extract of *A. esculentus* (Fig. 1**B**) are 3419 cm^−1^, corresponding to O–H stretching vibration, which indicates the presence of alcohol, 2921 cm^−1^ to C–H stretching of an aromatic compound, 2360 cm^−1^ to O–H stretching for carboxylic acid, 1647 cm^−1^ to C–C vibration and 1558 cm^−1^ to N–H stretching vibration present as the stabilizing and capping agents, as reported in a previous study (Kurian et al., 2022). The peak at 1384 cm^−1^ is attributed to C– O group, denoting carbonyl groups. The peak at 1033 cm^−1^ designates aliphatic amines’ C–N stretching vibration (Awwad and Salem, 2012). The FTIR results suggest that AgNPs were stabilized by terpenoid, alcohol, and carbonyl groups, providing strong binding sites for AgNPs. Thus, the FTIR results corroborate the presence of these biomolecules responsible for efficiently capping and stabilizing synthesized nanoparticles, which is consistent with previous studies (Zewde and Geremew, 2022)

#### 3.2.3 Energy dispersive X-ray (EDX) analysis

The EDX spectrum obtained from the green synthesized AgNPs showed a peak (∼3 keV) corresponding to elemental silver (Ag). The intensity of the EDX peak depends on the concentration of silver in the sample. The occurrence of the peak at 3 keV of EDX analysis is consistent with prior studies (Suba et al., 2022). Characteristic peaks of other elements like C and O, present in the sample, were also visible in the spectrum.

#### 3.2.4 X-ray diffraction (XRD) analysis

The crystalline phase and size of the synthesized AgNPs were determined using X-ray diffraction (XRD). The sample was placed on an XRD grid, and the diffraction patterns were recorded for the 2θ range of 20 to 80 degrees with a step of 0.0202 degrees (Bruker d8 Advance X-ray diffractometer, CuKα radiation, λ = 1.5406 Å, 40 kV-40 mA). XRD patterns revealed the presence of five major, distinct peaks at 38.33, 44.40, 64.53, and 77.49 Å, which correspond to the crystal planes (111), (200), (220), and (311) of silver, respectively. These peaks align with the powdered diffraction standard values of Miller indices (hkl) of the face-centered cubic (FCC) structure of silver (Fig. 3) and are consistent with the standard powder diffraction card of the Joint Committee on Powder Diffraction Standards (JCPDS) File No.: 04-0783 for silver (Gates-Rector and Blanton, 2019). The average size of the AgNPs was estimated using the Debye-Scherrer formula, D = 0.9λ/β Cos θ, where λ is the wavelength of the X-rays used for diffraction and β denotes the full width at half maximum (FWHM) of a peak. Based on the XRD spectrum of the AE-AgNP, the average AgNP size was 39.78 nm. A few unassigned peaks were detected, which may have been caused by the existence of bioorganic compounds or proteins in the extracts that crystallized on the surface of the AgNP (Sharma et al., 2022). In a similar observation, the AgNPs synthesized from the flower extract of *Mangifera indica* also showed a similar pattern of peaks in XRD analysis (Ameen et al., 2019).

**Figure 3.**
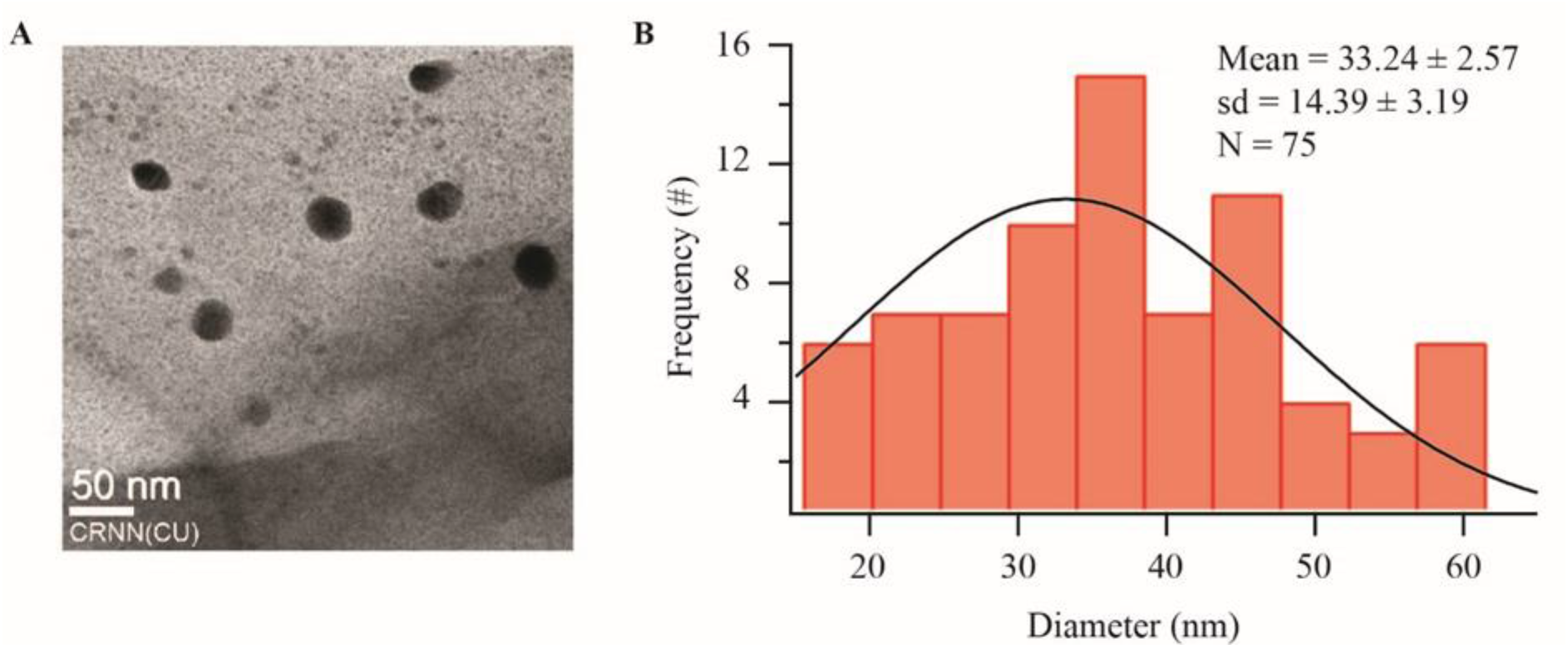
**A**. A typical TEM micrographic image of synthesized AgNPs. **B**. AgNPs size distribution extracted from TEM images. The solid black curve is a Gaussian fit to the data.

#### 3.2.5. Transmission electron microscopic (TEM) analysis

The particle size and crystalline state of the biosynthesized material were analyzed by TEM (JEM - 2100 HR, JEOL). Before the analysis, the sample was mixed with ethanol and ultrasonicated for 15 minutes to prepare a suspension which was then put on the copper grid and dried at 25°C (room temperature). Later it was positioned on a holder (specimen) for TEM analysis. The TEM images showed that the AgNPs were mostly spherical, with an average size of 33.24 ± 2.57 nm, consistent with the particle size range of AgNPs from *Ricinus communis* (Gul et al., 2021). A Gaussian data fit (Fig. 3B, black curve) to the histogram of the empirically measured particle size was performed to extract the average nanoparticle diameter and its spread due to the inhomogeneity of the nanoparticle sizes.

### 3.2 Efficacy of AE-AgNP on *Raillietina* spp

*Raillietina* spp. incubated in the control media (PBS alone) demonstrated physical activity for longer duration; the controls survived for approximately 72.00± 0.04h until becoming paralyzed and dead (Table 1; Fig. 4). When exposed to the test media (AE-AgNP and Genistein), the parasites transitioned from a strong movement activity to a relaxed state, then to paralysis and death. With dosages of 25, 50, 75, 100, and 125µg/ml PBS, respectively, the paralysis time was 1.57 h, 1.35 h, 1.06 h, 0.58 h, 0.54 h, and the death time was 2.54 h, 2.32 h, 1.54 h, 1.46 h, 1.29 h. A similar study showed that *Alpinia nigra*, a folklore plant used by the Tripuri tribe in North-East India, possesses significant anticestodal efficacy in a dose-dependent manner (Roy and Swargiary, 2009).

**Figure 4.**
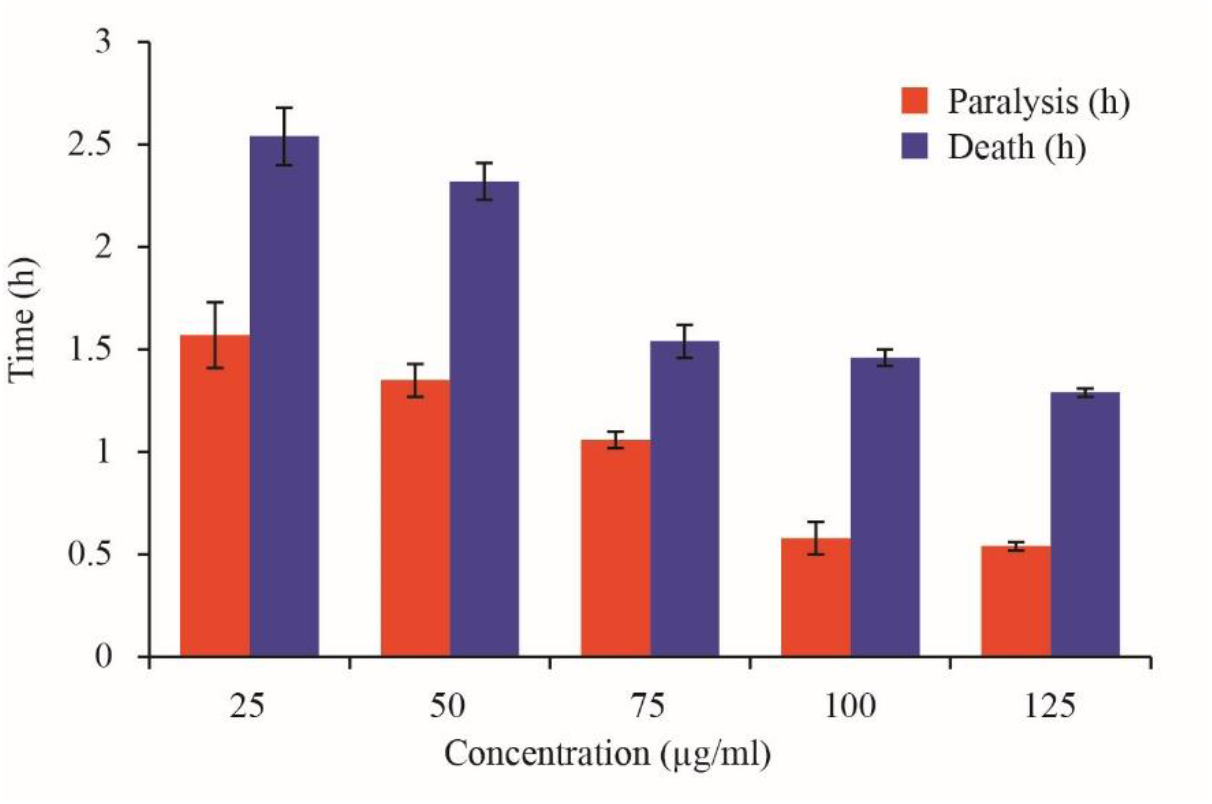
Results of AE-AgNP efficacy on *Raillietina* spp. after exposure to five different concentrations (25 µg/ml, 50 µg/ml, 75 µg/ml, 100 µg/ml and 125 µg/ml PBS).

**Figure 5.**
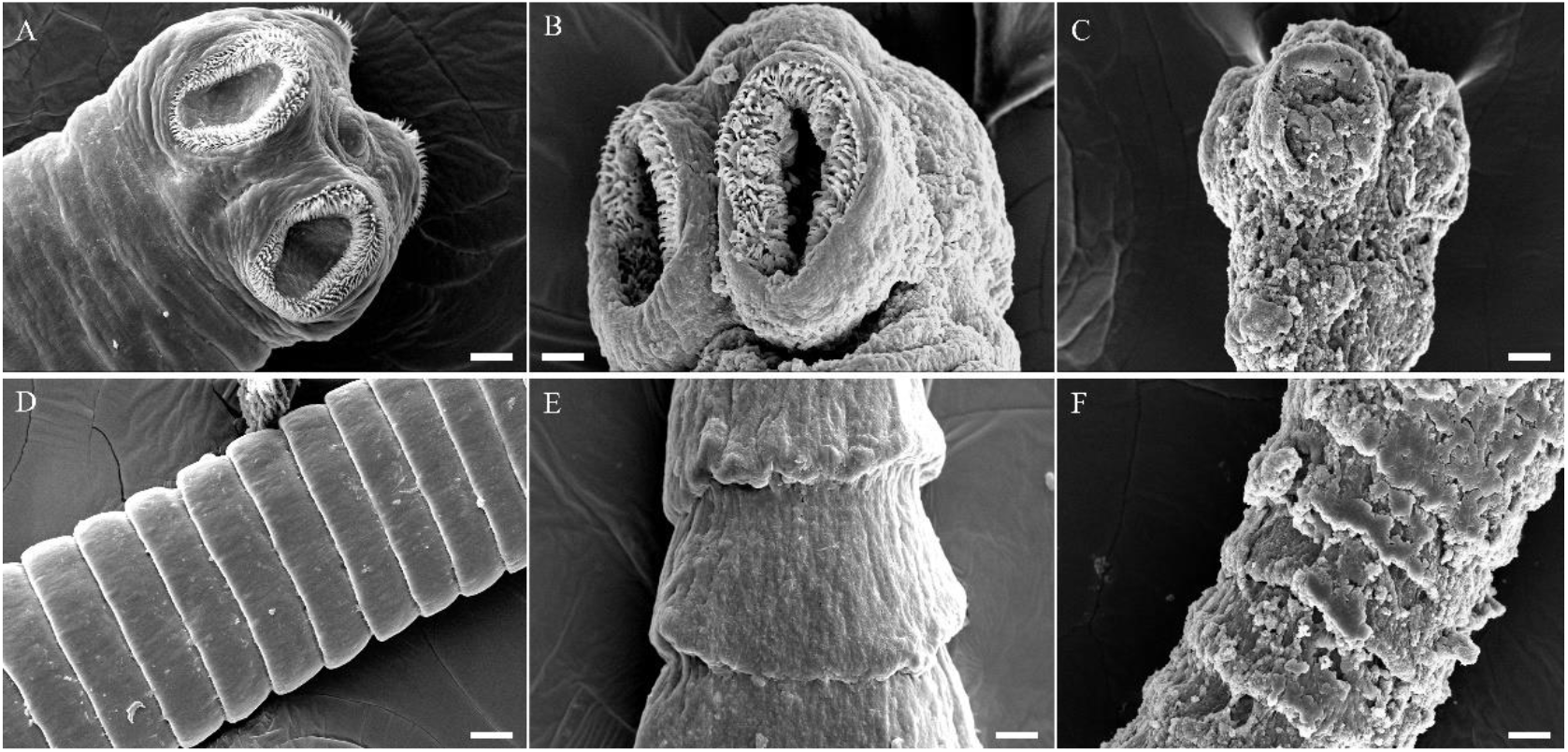
Scanning electron micrographs of control worm (**A** - scolex, **D** - gravid proglottid); Genistein (**B** - scolex, **E** - gravid proglottid) and silver nanoparticle exposed *Raillietina* spp. (**C** - scolex, **F** - gravid proglottid). All scale bars correspond to 20 μm.

### 3.3 Morphological changes of AE-AgNP exposed *Raillietina* spp

The *Raillietina* spp. is a long, segmented worm with distinct body regions, including the scolex (head), neck, and strobila (body proper). The scolex possesses four suckers and a rostellum, which are used for attachment. The proglottids, individual segments of the strobila, are covered in hair-like structures called microtriches that serve as the absorptive structures for feeding. Scanning electron microscopy (SEM) images show that the suckers on the scolex are arranged in a sideways pattern, with broad hooks at the base that tapers towards the end. The proglottids have a smooth, velvety appearance due to the unidirectional orientation of the microtriches covering their surface. However, when exposed to test media, the surface topography of the proglottids degenerated, resulting in the formation of wrinkles and erosion of the spines around the suckers, which altered the host-parasite interface. Treatment with Genistein, a reference drug, caused significant damage to the scolex and the tegumental surface structures, leading to their breakage and detachment. In a similar study, on incubating the cestodes with the root-peel extract of *Potentilla fulgens*, the parasites exhibited complete attrition of microtriches from the tegument, disintegration of muscle bundles and cellular organelles (Roy et al., 2012).

### 3.4 Histochemical studies

The activity of AcPase, AlkPase, ATPase, and 5’-Nu was most intense in the tegument (T) of control *Raillietina* spp. as compared to the sub-tegument (ST) and somatic musculature (SM), as shown in Figs. 6**A**-**D**. The parasites exposed to AE-AgNP revealed a general reduction in staining intensity in the T, ST, and SM, while no activity was visible in parenchyma cells (P), as seen in Figs. 6**E**-**H**. Specifically, the staining intensity of AcPase was significantly reduced in the T and ST regions of the AgNP- incubated section (Figs. 6**A, E**), whereas the activity throughout the section of parasites exposed to Genistein was minimal (Figs. 6**I**-**L**). AlkPase activity was also greatly diminished throughout the treated sections of the parasite (Figs. 6**B, F**), and ATPase activity was almost imperceptible in the parasite’s T, ST, SM and P treated with AE-AgNP compared to control parasites (Figs. 6**C, G**). Finally, 5’-Nu activity was also reduced along the T, ST and SM region in the AgNP-exposed cestodes compared to the control (Figs. 6**D, H**). Histochemical studies have evaluated the parasite’s localization and expression of specific enzymes and metabolic pathways in response to phytochemical treatment. Another study revealed potential antitrematocidal activity by *Senna* leaf extracts against *Paramphistomum gracile* by altering tegument architecture and inhibiting tegumental enzyme activity (Roy and Lyndem, 2019). Phytocompounds such as *α*-viniferin altered the tegumental morphology of the treated parasites, reducing the tegumental enzyme’s activities (Roy and Giri, 2015). Another study observed a decrease in the levels of phosphatases and trace elements on exposure to crude ethanol extract of ethnomedicinal plants *Acacia oxyphylla* and *Securinega virosa*, compared to control groups (Dasgupta et al., 2013).

**Figure 6.**
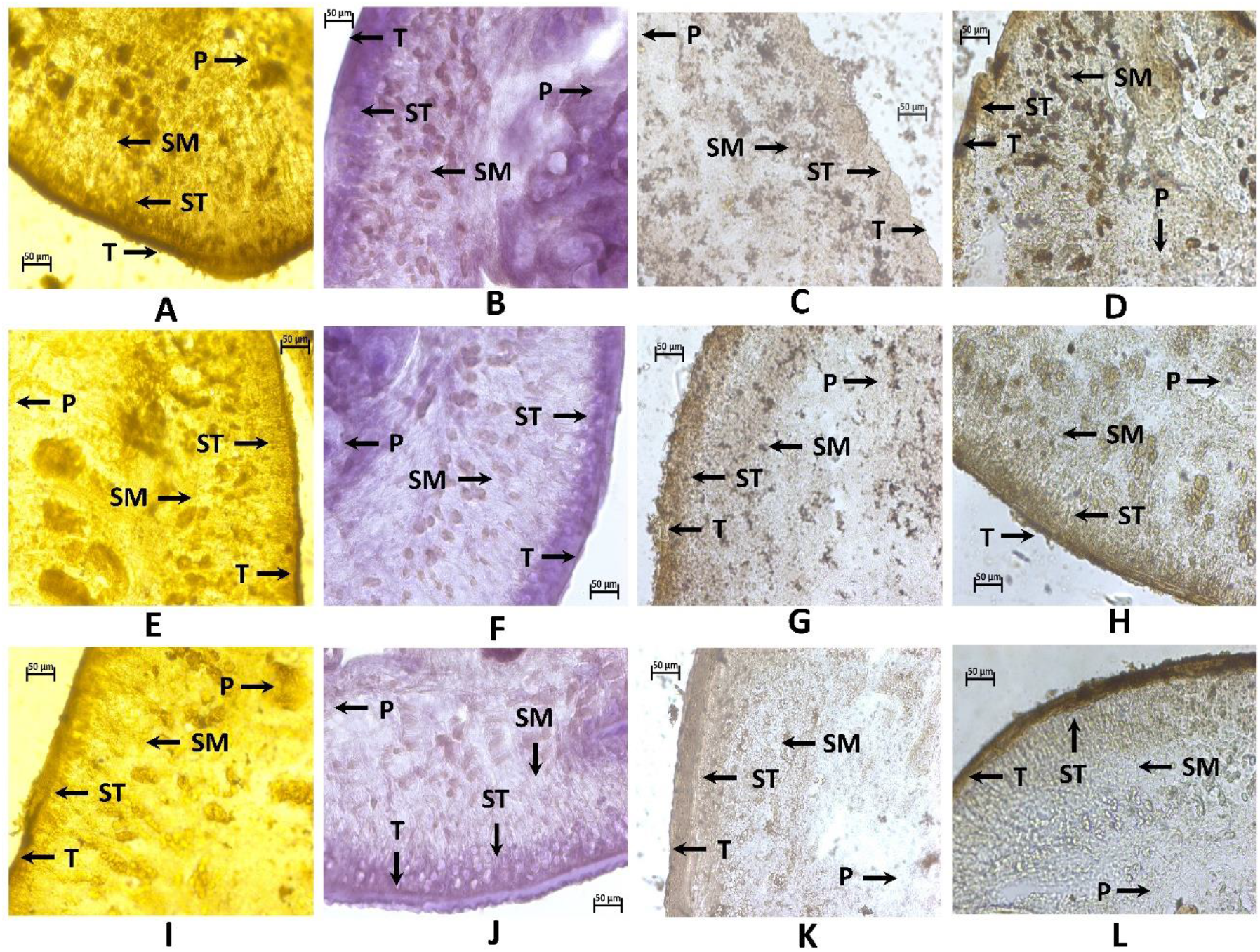
Histochemical demonstration of AcPase (**A, E, I**), AlkPase (**B, F, J**), ATPase (**C, G, K**) and 5’-Nu (**D, H, L**) activities in *Raillietina* spp. treated with AE-AgNP (125 μg/ml) and Genistein GEN (125 μg/ml); **A**-**D**: Transverse section of control parasite; **E**-**H**: AgNPs -exposed parasite; **I**-**L**: GEN- exposed parasite. All scale bars correspond to 50 μm.

## Conclusion

The study describes a green synthesis method for AgNPs using an extract from *Abelmoschus esculentus*. The biosynthesized AgNPs were spherical, had a 30-35 nm size range, and showed significant anthelmintic activity against the model cestode. Electron microscopy and histochemical studies have provided important insights into the mechanism of action of phytochemicals and helped identify potential targets for anthelmintic therapy. These studies have provided important implications for developing alternative and more sustainable approaches to controlling parasitic cestode infections in poultry. Green synthesized anthelmintics can be safer and more effective and are less likely to have harmful side effects or produce resistant strains of helminths. The plant extracts used in green synthesis have cultural significance in ethnomedicinal practices, and using these extracts to synthesize anthelmintics can help preserve traditional knowledge and promote cultural diversity. This study illustrates the possibility of using plant derivatives for the biosynthesis of AgNPs and enunciates the need to develop environmentally benign methods for synthesizing nanomaterials.

## Authorship contribution statement

Rima Majumdar: Writing - original draft, conceptualization, investigation and data analysis. Pradip Kumar Kar: Experiment - designing, Writing - review and editing, supervision.

## Acknowledgments

The authors thank Dr. Debkumar Mukhopadhyay, Vice-Chancellor of CBPBU, for his valuable support, encouragement, and provision of essential laboratory facilities. Furthermore, the authors thank the UGC-DAE Consortium in Kolkata for providing the XRD facility and the Centre of Nanoscience and Nanotechnology (CRNN) in Kolkata for granting permission to utilize their SEM, TEM, and FTIR facility.

